# EVALUATION OF STED SUPER-RESOLUTION IMAGE QUALITY BY IMAGE CORRELATION SPECTROSCOPY (QuICS)

**DOI:** 10.1101/2021.08.30.457899

**Authors:** Elena Cerutti, Morgana D’Amico, Isotta Cainero, Gaetano Ivan Dellino, Mario Faretta, Giuseppe Vicidomini, Pier Giuseppe Pelicci, Paolo Bianchini, Alberto Diaspro, Luca Lanzanò

## Abstract

Quantifying the imaging performances in an unbiased way is of outmost importance in super-resolution microscopy. Here, we describe an algorithm based on image correlation spectroscopy (ICS) that can be used to assess the quality of super-resolution images. The algorithm is based on the calculation of an autocorrelation function and provides three different parameters: the width of the autocorrelation function, related to the spatial resolution; the brightness, related to the image contrast; the relative noise variance, related to the signal-to-noise ratio of the image. We use this algorithm to evaluate the quality of stimulated emission depletion (STED) images of DNA replication foci in U937 cells acquired under different imaging conditions. Increasing the STED power improves the resolution but may reduce the image contrast. Increasing the number of line averages improves the signal-to-noise ratio but facilitates the onset of photobleaching and subsequent reduction of the image contrast. Finally, we evaluate the performances of two different separation of photons by lifetime tuning (SPLIT) approaches: the method of tunable STED power and the commercially available Leica Tau-STED. We find that SPLIT provides an efficient way to improve the resolution and contrast in STED microscopy.

## INTRODUCTION

Super-Resolution Microscopy (SRM) circumvents the spatial resolution limit imposed by the diffraction of light at about half of the illumination wavelength (200-250 nm for visible wavelength). Among the super-resolution techniques developed in the last decade, some of them, grouped under the term of nanoscopy, can theoretically reach diffraction-unlimited resolution, down to molecular size ^1^. The common working principle of these techniques is to transiently transfer the fluorophores in two recognizable states (usually a dark OFF state and a bright ON state in response to different stimuli); this allows the subsequent sequential detection of signals originating from regions much smaller than the diffraction limit ^2, 3^. Among the super resolution techniques that do not require a complex image reconstruction process, the most used is Stimulated Emission Depletion microscopy (STED): STED microscopy overcomes the diffraction limit by reversibly switching off (depleting) fluorophores at the periphery of the diffraction-limited excitation regions. The depletion is achieved thanks to a second beam (the so-called STED beam) tuned in wavelength to induce stimulated emission and engineered in phase to create a doughnut-like shaped intensity profile at the focus. By increasing the intensity of the STED beam, stimulated emission wins the competition against spontaneous emission of fluorophores and allows to register fluorescence only from those fluorophores localized in a tiny sub-diffraction volume at the center of the excited region ^4^.

Although nanoscopy techniques and STED, in particular, can theoretically achieve unlimited resolution, experimental constraints on biological samples considerably reduce the spatial resolution improvement to about 20 nm. Moreover, a series of factors related to cell labelling ^5, 6^ and image acquisition ^7–11^ must be carefully assessed and adjusted depending on the biological mechanism under investigation. Examples of acquisition parameters that must be carefully adjusted in STED microscopy are the STED beam intensity, the excitation beam integration and the pixel dwell-time. Typically, one has to find a trade-off between several conditions to avoid the onset of unwanted sample degradation effects such as fluorophore photobleaching. This trade-off is often specific for the biological sample considered and cannot be easily determined using calibration samples (i.e., fluorescent spheres). Thus, quantifying the imaging performances directly on the acquired images, in an unbiased way, is of outmost importance ^10, 12–15^.

Here, we introduce a simple algorithm to evaluate systematically and in an unbiased way the quality of STED images by image correlation spectroscopy (QuICS). Image correlation spectroscopy (ICS) is a general and versatile method to quantitatively analyze fluorophore distribution in microscopy images ^16^. ICS can be used to extract parameters such as size ^17^, distances ^18, 19^ and aggregation state ^20^ from static images and dynamic parameters such as diffusion coefficient ^21^ and velocity ^22^ from time-resolved images. In this work, we focus only on the analysis of static images. We apply ICS to extract three quantities that are related to the quality of the super-resolved image: the width of the autocorrelation function, related to the spatial resolution; the brightness, related to the image contrast; the relative noise variance, related to the signal-to-noise ratio of the image. Within this study, we describe how the modulation of image acquisition parameters can influence STED efficiency and the image formation of DNA replication sites in U937-PR9 cells, an in vitro model of leukemia ^23^. Our study reveals that to optimize the imaging conditions for a given sample, a balance between different parameters must be found. We found a valid solution to this elusive balance by applying the method of Separation of Photons by LIfetime Tuning (SPLIT) ^24, 25^ to STED microscopy. In particular, we show SPLIT images obtained using the method of tunable STED power ^25, 26^, or acquired by a recently developed, lifetime-based commercial setup (the Leica Tau-STED microscope). QuICS analysis reveals that SPLIT images have higher resolution and non-reduced brightness and noise parameters, compared to their counterpart STED images.

We developed the QuICS algorithm base on the growing need for analysis of nuclear processes performed at the level of individual cells, also taking into account that certain events typically occur in a relatively small fraction of cells in the population at any given time (i.e., events taking place in a specific phase of the cell cycle). Recent advances observed a considerable variability and heterogeneity in genome organization at the single-cell level ^27^. Imaging and super-resolution can thus provide a unique view of nuclear organization and functions in intact cell nuclei.

## RESULTS

### Autocorrelation function as a source of information about image quality

In order to extract a series of parameters associated to an image I(x,y), we start by calculating a radial autocorrelation function (ACF) G(ρ). This function is calculated by performing an angular average on the two-dimensional ACF G(δx,δy) (see Methods). In general, the function G(ρ) contains information on all the intensity variations in the image, including fluctuations due to statistical noise. Let’s call G_NF_(ρ) the noise-free correlation function, i.e. the corresponding function in the absence of noise. By fitting G_NF_(ρ) to a Gaussian model (see Eq. (5)), we extract the amplitude G_NF_(0) and the width parameter w. We define the following three quantities (Figure 1):

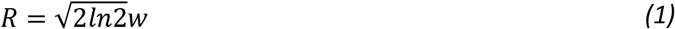

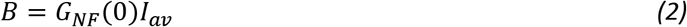

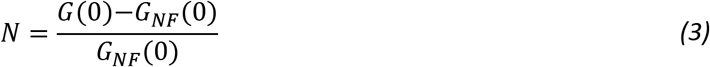

**Figure 1.**
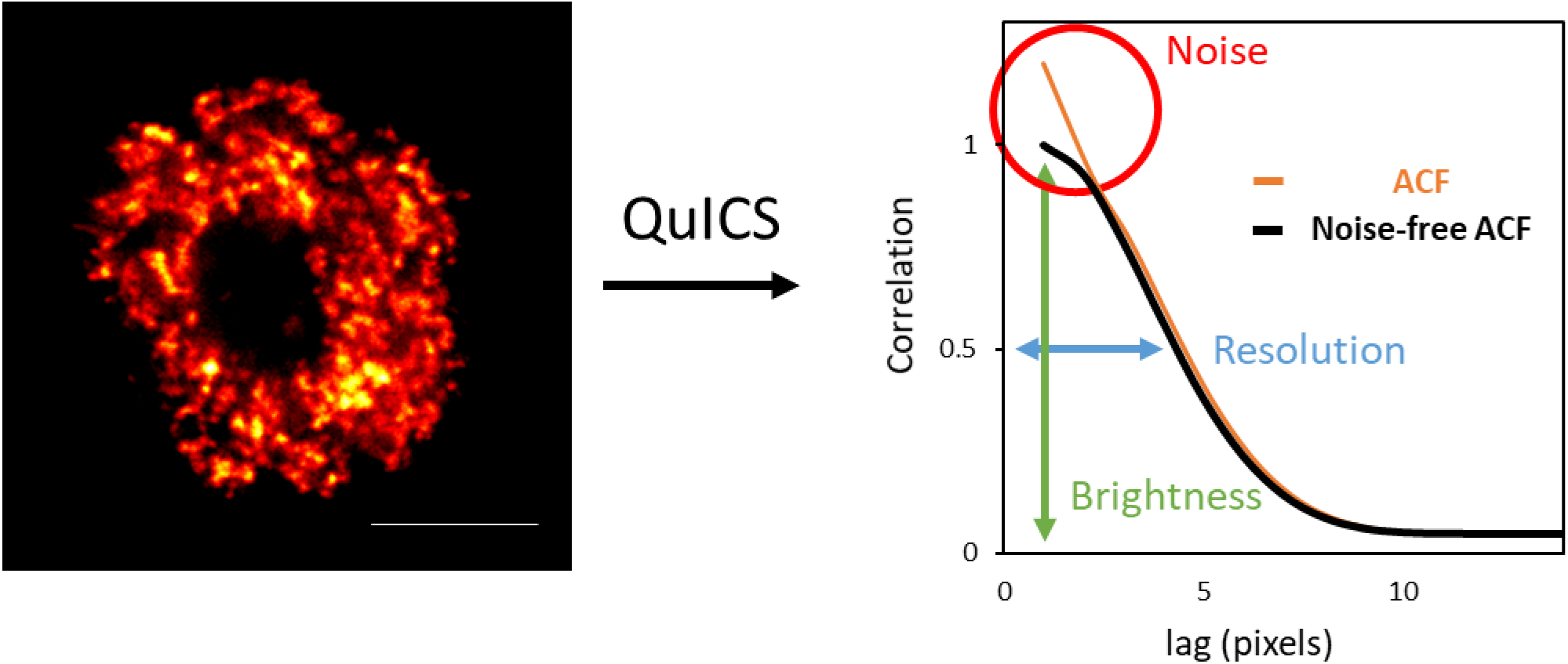
Autocorrelation function (ACF) as a source of information about image quality. Schematic representation of the application of the QuICS algorithm to an image of a nuclear process (DNA replication sites). The algorithm calculates a radial autocorrelation function (ACF, orange line) and performs a Gaussian fit of the estimated noise-free ACF (black line). The three parameters that are extracted are: the Resolution (in blue), calculated from the width of the noise-free ACF, the Brightness (in green), calculated from the amplitude of the noise-free ACF, and the Noise (in red), calculated from the difference in amplitude between the ACF and the noise-free ACF. Scale bar represents 3 μm.

Where we have indicated I_av_ as the average intensity value over all the pixels of the image.

In order to understand the physical meaning of R, B and N, let’s assume, for simplicity, that the sample contains randomly distributed point-like fluorescent particles so that the corresponding image is the convolution of the emitters and the Point Spread Function (PSF) of the optical system. In this case, R corresponds to the resolution of the optical system expressed in terms of the Full Width at Half Maximum (FWHM) of the PSF, R=FWHM_PSF_. More in general, since the sample may contain features of finite size, it will be R≥FWHM_PSF_. Thus, the estimated resolution of the optical system is at least equal to R.

The quantity B in Eq. (2) is called brightness ^28, 29^ and is equal to σ_p_^2^/I_av_ where σ_p_^2^ is the variance of the intensity due to the particles. The brightness of the particles depends on the number of fluorophores per particle and on the actual brightness of the fluorophores at the specific imaging settings (e.g. excitation intensity level, detector gain, pixel dwell time). Let’s assume that, in addition to the signal from the particles, whose average intensity is I_av,p_, there is a uniform background signal, with average intensity I_av,bkgd_, so that I_av_= I_av,p_ + I_av,bkgd_. The brightness is given by B= σ_p_^2^/(I_av,p_ + I_av,bkgd_). Thus, a reduction of the brightness parameter B is related to a decrease in the contrast of the particles in the image.

Finally, we note that G_NF_(0)=σ_p_^2^/I_av_^2^ and G(0)= (σ_p_^2^+ σ_noise_^2^)/I_av_^2^, where σ_noise_^2^ is the variance of the intensity due to noise. Thus, the quantity N in Eq. (3), N=σ_noise_^2^/σ_p_^2^, represents the variance of the noise normalized to the variance of the particles (relative noise variance). We use N to quantify the noise level in the image, where the limits N=0 indicates no noise, and N=1 indicates that intensity fluctuations due to noise are comparable to those due to the particles.

The noise-free correlation function, G_NF_(ρ), required for this analysis, can be obtained by cross-correlating two statistically independent realizations of the same image, in analogy to what is done in Fourier Ring Correlation (FRC) methods ^12–14^. The acquisition of two statistically independent images is straightforward in single-molecule localization microscopy but can be a more difficult task with other super-resolution techniques ^15^. In order to estimate G_NF_(ρ) from a single acquired image, we propose performing a fit of the autocorrelation function G(ρ) either skipping the first points, or cross-correlating two statistically independent images obtained by down-sampling the original image ^30^. In our experimental data, the two approaches provided similar results (Supplementary Figure 1).

### Tuning of STED power

During STED image acquisition, the most intuitive way to improve the resolution is to increase the STED beam’s intensity. To validate our method as a function of the STED beam intensity, we acquired images of U937-PR9 cells samples in which we were able to visualize the DNA replication thanks to the incorporation of the nucleoside Ethynyl deoxyuridine (EdU) to the newly replicated DNA strand. We then coupled the EdU molecule to an azide molecule carrying the Alexa fluorophore, taking advantage of a Cu-catalyzed Click-iT reaction. During microscopy acquisition, we took care of choosing, among the sample, actively replicating cells with DNA replication sites spread all over the nucleus and thus more suitable for resolution evaluation analysis.

First, we acquired a confocal and a STED image of a cell nucleus (Figure 2A, upper row) by applying a relatively small depletion beam power (9 mW) and the two line profiles of the same structure were plotted to compare the achieved resolution (Figure 2B, upper panel). Then, we acquired a confocal and a STED image of a second nucleus, doubling the power of the depletion beam (Figure 2A, lower row), and we compared the line profiles of the same structure from the confocal and the STED image. The line profile analysis yielded FWHM=189 nm and FWHM=212 nm for the two peaks detected at 9 mW, and FWHM=160 nm and FWHM=178 nm for the two peaks detected at 18 mW. However, this result depends on the specific structures selected for the line profile analysis and does not take into account the totality of labeled sites in the whole cell nucleus.

**Figure 2.**
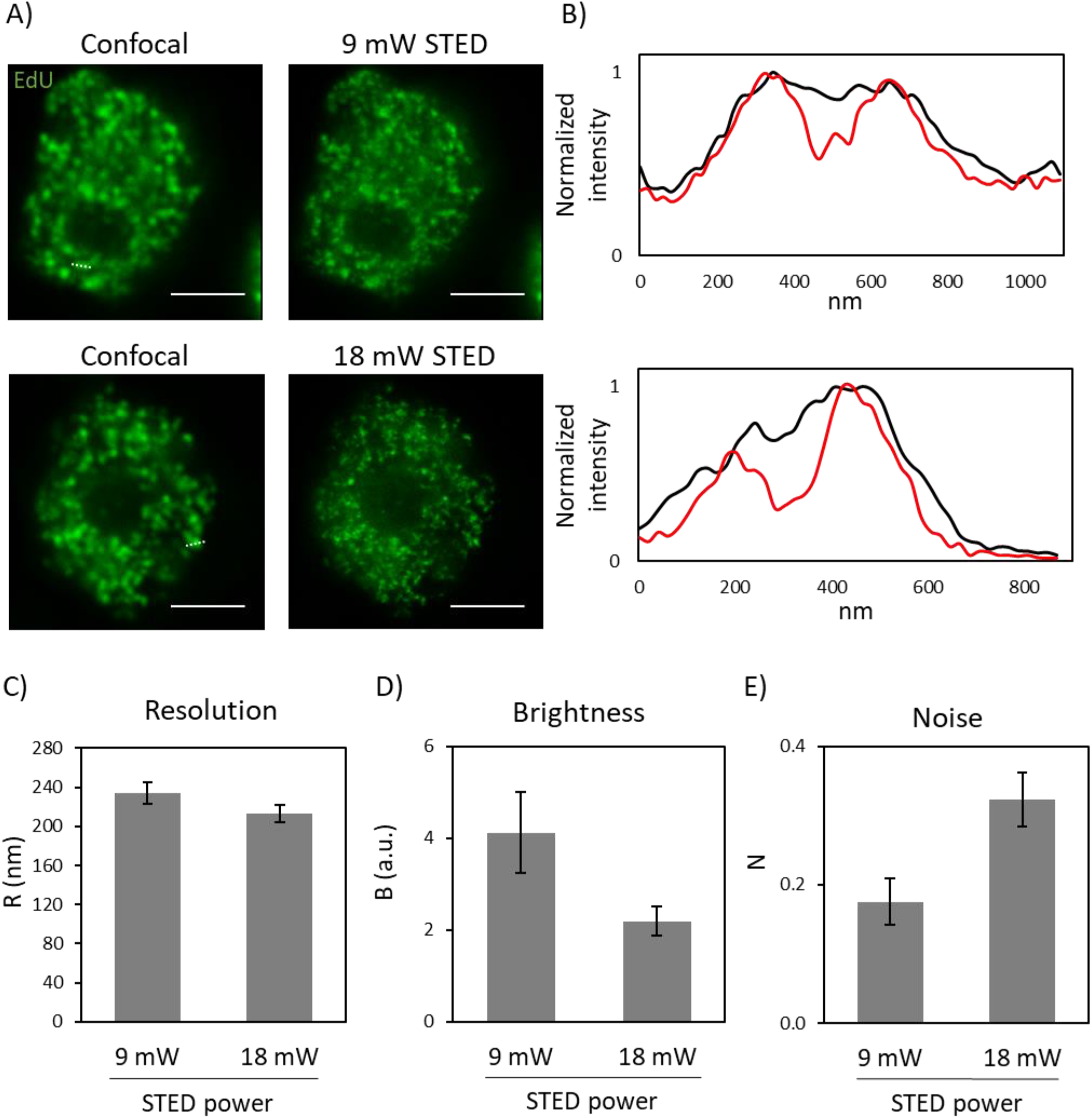
Tuning of STED power. **A)** Representative images of U937-PR9 cells upon staining of DNA replication foci through incorporation of EdU labeled with Alexa azide 488 (Click reaction). (top) Sequential acquisition of a confocal image, followed by a STED image with a 9 mW depletion beam. (bottom) Sequential acquisition of a confocal image, followed by a STED image with a 18 mW depletion beam. Scale bars represent 3 μm. **B)** Line profiles of structures from images in A). (top) Comparison between the line profiles of the same structure in the confocal (black line) and the STED (red line) images in the top row of panel A). The measured structure is defined by a white dotted line in the confocal image. (bottom) Comparison between the line profiles of the same structure in the confocal (black line) and the STED (red line) images in the bottom row of panel A). The measured structure is defined by a white dotted line in the confocal image. **C) D) E)** Quantification of Resolution, Brightness, and Noise parameters by application of the QuICS algorithm. At least ten images for each condition have been acquired. Error bars represent the SEM.

Therefore, we acquired at least ten STED images with each of the two different depletion beam powers, and we calculated the autocorrelation function in order to obtain the average resolution R related to the entire nuclei. As a result, we obtained that the doubling of the STED beam lead to an improvement of spatial resolution from R=234 ± 3 nm to R=213 ± 3 nm (mean ± s.d., Figure 2C). This result is in keeping with the line profile analysis, as expected, and represents an average of the whole nucleus structures. The obtained values of R strongly depend on the average apparent size of the structures (i.e. replication foci) in the images, meaning the molecular volume plus the resolution, and therefore their values are larger than the maximum resolution of the optical system.

In contrast, we observed that the image brightness B was significantly lower for the images acquired with the higher STED power (Figure 2D). We interpreted this reduction of B as a reduction in the image contrast. In fact, we compared images acquired exactly with all the same instrumental settings (e.g. same excitation power, same detector gain, same pixel dwell-time) other than the STED power. In this case, the action of STED reduces the average intensity per pixel due to the particles but does not decrease the average intensity of the background signal (originating, for instance, from undepleted out-of-focus fluorescence signal ^24, 31, 32^). As explained in the previous section, this causes a reduction of the parameter B.

Finally, we observed that the relative noise variance N was higher at the higher STED power (Figure 2E). This is in line with the expected reduction of signal-to-noise ratio at increasing STED power.

### Increasing number of averages

The common approach for reducing the noise in an image is to increase the number of collected photons per pixel. In general, this can be achieved by tuning the number of scans for each pixel and then averaging the intensity values for each pixel position.

To evaluate how increasing averages would influence the quality of the image, we acquired sequential STED images of the same cell, with a depletion beam power of 18 mW (see Methods for a detailed description of the sequential acquisition settings). From each sequential acquisition, we generated STED images with a different number of line-averages (Figure 3A) and applied the QuICS algorithm. As expected, the image’s noise significantly decreased as soon as we doubled the number of line averaging (Figure 3B). On the other hand, the average resolution R did not improve with an increasing number of averages (Figure 3C). The brightness B decreased as a function of the number of averages (Figure 3D) indicating a reduction of the contrast.

**Figure 3.**
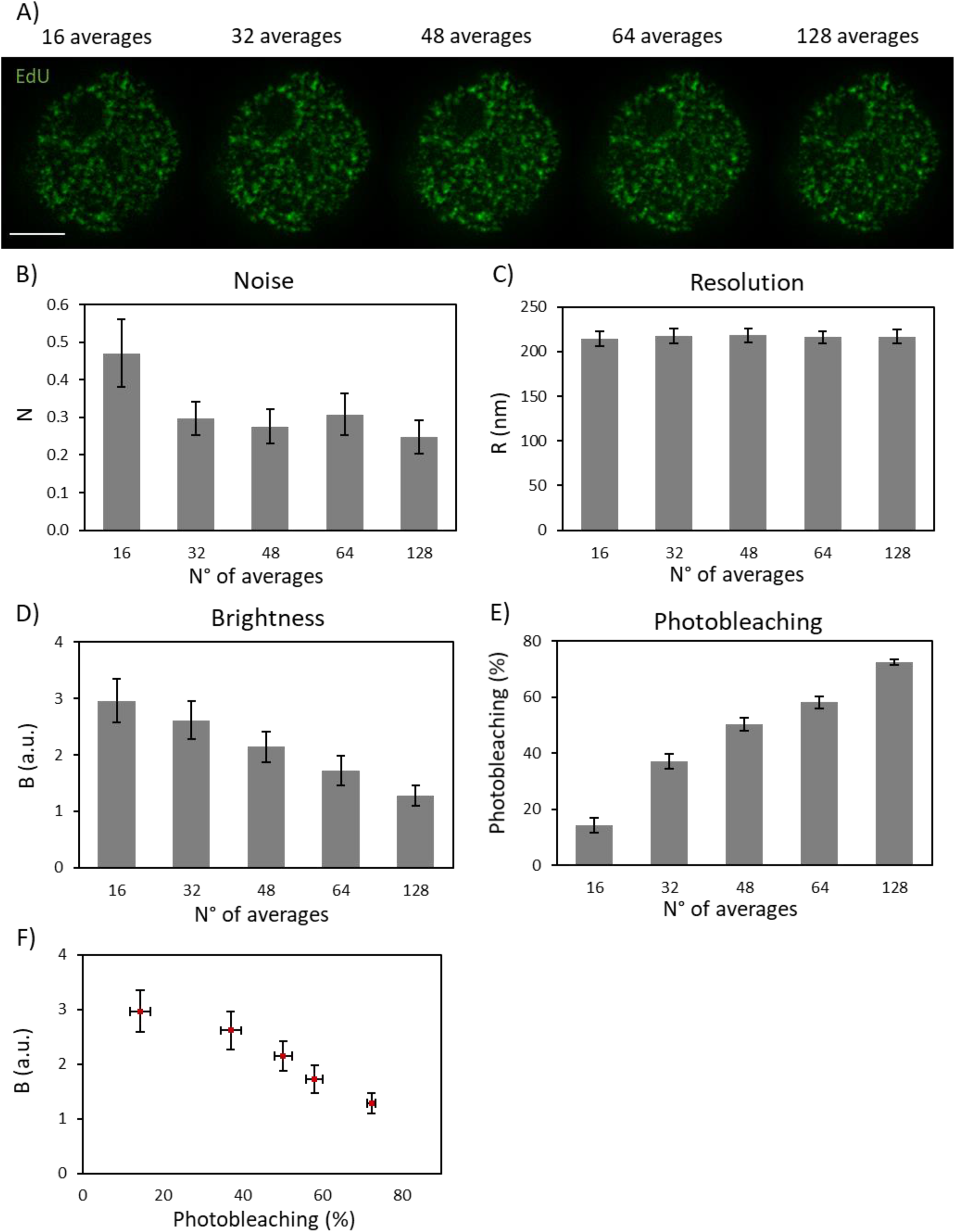
Increasing number of averages. **A)** Representative images of U937-PR9 cells upon staining of DNA replication foci. Each image is obtained by combining and averaging STED images after each acquisition step of increasing number of line-averages (see Methods for a detailed description of acquisition settings) Scale bar represents 3 μm. **B) C) D)** Quantification of Resolution, Brightness and Noise parameters in function of the number of line-averages by application of the QuICS algorithm. At least ten images for each condition have been acquired. **E)** Quantification of Photobleaching in function of the number of line-averages. Photobleaching was calculated as the percentage reduction of average fluorescence intensity with respect to the initial value. **F)** Representation of the Brightness variation in function of the Photobleaching. At least ten images for each condition have been quantified. Error bars represent the SEM.

To interpret these results, we monitored photobleaching as a function of the number of averages of the STED image (Figure 3E). Photobleaching was calculated as the percentage reduction of average fluorescence intensity with respect to the initial value. We observed that each line averages-acquisition step induced a significant increase in photobleaching of the sample’s fluorophores (Figure 3E). Consequently, the image contrast decreased, thus leading to a brightness reduction as a function of the number of averages (Figure 3D,F). These data also show that, in our samples, for photobleaching levels above about 40%, there is no significant improvement in the signal-to-noise ratio of the images (Figure 3B,F). Thus, in case it cannot be avoided, photobleaching should be at least kept below this level.

### Comparison between SPLIT and STED imaging

To increase the spatial resolution of a STED microscope, the most straightforward way is to increase the depletion beam’s intensity. However, as we have seen, this may reduce the contrast and signal-to-noise of the images, quantified in QuICS via the brightness and noise parameters. Here, we examine the advantages of increasing spatial resolution via application of Separation of Photons by LIfetime Tuning (SPLIT) ^24^. The SPLIT method provides an increase in spatial resolution by decoding the spatial information encoded into an additional channel. The first reported SPLIT configuration exploited, as an additional channel, the fluorescence lifetime gradient induced by a continuous-wave STED beam ^24, 33^. Subsequent studies demonstrated that SPLIT is not limited to analysis of fluorescence lifetimes. SPLIT could also be applied to stacks of STED images obtained with tunable depletion power, with the depletion power used as the additional channel for SPLIT ^25, 26^, or even to structured illumination microscopy images ^34^.

As described in Figure 4A, we first applied the SPLIT method to stacks consisting of two STED images at different depletion power: a confocal (0 mW STED power) and a STED image (18 mW STED power). In this case, the fluorescence intensity variations due to the tuning of the STED power, allow the separation of the contributions from fluorophores in the center or the periphery of the PSF (Figure 4A). Since the excitation intensity can also be easily tuned along the stack ^26^, we set the excitation level of the confocal image so that it induced negligible photobleaching. In this way, the data acquisition for SPLIT was straightforward and did not induce more photobleaching than the acquisition of the STED image alone. Figure 4B shows application of this approach to imaging of replication foci in a U937-PR9 cell in S phase. Shown are the confocal, the STED image and the resulting SPLIT image. We compared the line profiles of the same structure and we observed a resolution improvement from FWHM=147 nm and FWHM=154 nm, for the two peaks detected in the STED line profile, to FWHM=135 nm and FWHM=107 nm, for the two peaks detected in the SPLIT line profile (Figure 4C). QuICS analysis of at least ten samples, revealed a significant improvement in the average resolution of the SPLIT image (R=129 ± 9 nm, mean ± s.d.) compared to the STED image (R=213 ± 9 nm, mean ± s.d.) (Figure 4D). Notably, this improvement in resolution is not achieved at the expense of the image brightness (Figure 4E) or the signal-to-noise ratio (Figure 4F).

**Figure 4.**
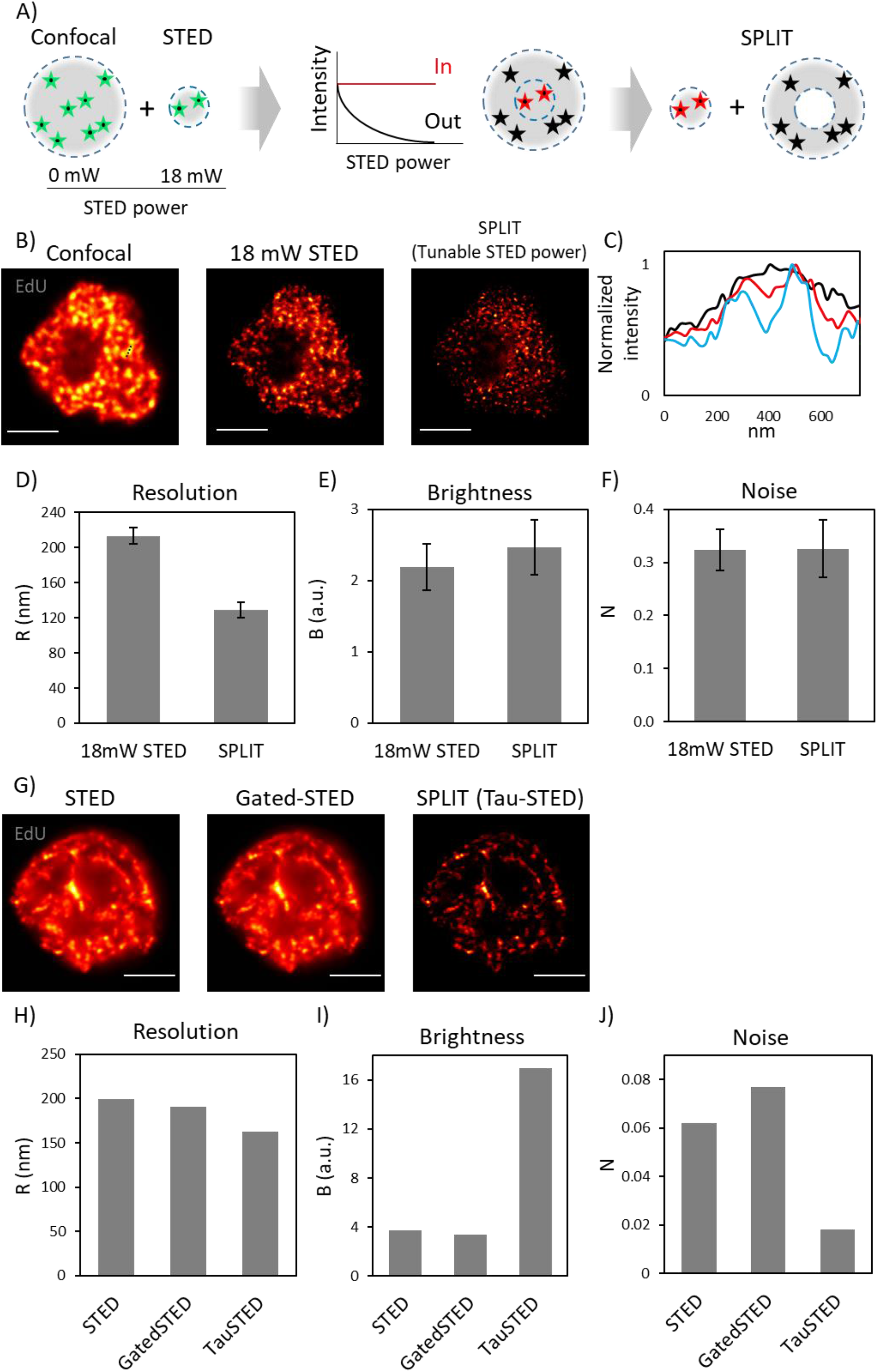
Comparison between SPLIT and STED imaging. **A)** Schematic representation of the SPLIT principle using a tunable STED power. The sequential acquisition with an increased STED power from 0 to 18 mW, allow to obtain the extra information about the fluorescence depletion dynamics of photons arising from the center (in) and the periphery (out) of the PSF. **B)** Representative images of U937-PR9 cells upon staining of DNA replication foci. Sequential acquisition of a confocal image (STED power: 0 mW) and a STED image with a 18 mW depletion beam, followed by the resulting SPLIT image. Scale bars represent 3 μm. **C)** Comparison between the line profiles of the same structure in the confocal (black line), the STED (red line) and the SPLIT (blue line) images of panel B). The measured structure is defined by a black dotted line in the confocal image. **D) E) F)** Quantification of Resolution, Brightness, and Noise parameters by application of the QuICS algorithm. At least ten 18 mW STED images have been acquired and compared to the resulting SPLIT images. Error bars represent the SEM. **G)** Images of U937-PR9 cells upon staining of DNA replication foci through incorporation of EdU labeled with Alexa azide 594. Images acquired with the Leica Stellaris 8 Tau-STED microscope. Shown are the raw STED image, the gated-STED image with a time-gating of 1-8 ns and the Tau STED image. Scale bars represent 3 μm. **H) I) J)** Quantification of Resolution, Brightness, and Noise parameters of images shown in G) by application of the QuICS algorithm. The analysis in G-J is representative of the analysis on three cells yielding similar results.

Figure 4G shows an image of replication foci in a U937-PR9 cell in late S-phase acquired with the Leica Tau-STED microscope. Here the SPLIT image (i.e. the Tau-STED image) is compared with the STED image and with a time-gated STED image (time gate=1-8 ns). QuICS analysis indicates an improvement of resolution from R=199 nm (STED image) and R=191 nm (gated-STED image) to R=163 nm (Tau-STED image) (Figure 4H). The brightness is significantly higher in the Tau-STED image than in the STED and gated-STED images (Figure 4I). This increase in brightness is probably due to the improvement of contrast provided by SPLIT, which has the capability of filtering out background signal originating for instance by direct excitation from the STED beam ^24^. The gated-STED image has lower SNR than the STED image, as time-gating reduces the number of photons available for image formation (Figure 4J). The SPLIT image has higher SNR than the STED and gated-STED images, in line with the overall reduction of background in the image (Figure 4J) and in keeping with previous studies ^35^.

## DISCUSSION

Applications of super-resolution microscopy to biology are increasing. However, despite the availability of several types of commercial setups, optimization of the conditions of imaging still requires some degree of expertise. It is important to find the conditions that maximize the quality of the image, paying attention to the onset of potentially degrading effects such as fluorophore photobleaching. Our approach provides an unbiased measurement of the super-resolution image quality based on the three parameters R, B, N defined by Eq. (1)(2)(3). We note that R and N can be readily used to compare the resolution and signal-to-noise ratio of images acquired under different conditions. On the other hand, B depends on the image contrast, but also on many instrumental factors (e.g., the excitation intensity level, the detector gain, and the pixel dwell-time) that should be considered when performing any comparison.

There is an important difference between QuICS and the FRC method. The FRC resolution merges into a single parameter information about both the relevant spatial frequencies and the noise content of an image. In other words, the FRC resolution describes the length scale below which the image lacks signal content ^13^. In QuICS, the resolution parameter R contains average information on the characteristic size (e.g. specimen features, PSF of the optical system) whereas the parameter N contains information on the noise content of the image. In the limit of infinitely high signal-to-noise ratio, the two values of resolution extracted by QuICS and FRC are the same. Conversely, for low signal-to-noise ratio, we expect the FRC value to increase whereas the QuICS resolution to remain constant, since it represents the average apparent size of particles in the image (or the size of the PSF, in the limit of point-like particles). Thus, an advantage of QuICS is that the same algorithm can be used not only to evaluate the image quality but also to quantify biophysical parameters such as the size and the molecular brightness, important in many biophysical applications ^16, 17, 28, 29, 36^.

The combination of super-resolution microscopy with the correlation spectroscopy toolbox undoubtedly offers several advantages ^37^. Here, we have shown how analysis of an angle-averaged, image correlation function can provide useful hints on the optimization of the imaging conditions. As a case study, we have focused our attention on STED imaging of DNA replication foci in fixed U937 cells. However, even if not demonstrated, we expect that the approach can be adapted to images containing arbitrary features (for instance cytoskeletal structures). Similarly, we expect that it can be used to evaluate the quality of images acquired in confocal microscopy or other types of super-resolution techniques.

## METHODS

### Cell culture and treatments

U937-PR9 cells were cultured in RPMI-1640 medium (Sigma Aldrich R7388) supplemented with 1% penicillin/streptomycin (Sigma-Aldrich P4333) and 10% fetal bovine serum (Sigma-Aldrich F9665) and maintained at 37°C and 5% CO_2_. U937-PR9 were seeded on poly-L-lysine (Sigma-Aldrich P8920) coated glass coverslips immediately before experiments. Cells were incubated with 10 μM of the synthetic nucleoside 5-Ethynyl-2’-deoxyuridine (EdU) (Thermo Fisher Scientific) for 25 min at 37°C and 5% CO_2_.

### EdU fluorescent labelling

Upon nucleoside incorporation, cells were washed with Phosphate Buffer Saline (PBS), fixed with 4% paraformaldehyde (w/v) for 10 min at room temperature and permeabilized with 0.5% (v/v) Triton X-100 in PBS for 20 min. Cells were then incubated for 30 min with the Click-iT reaction cocktail containing Alexa Fluor azide 488 (Invitrogen C10337) or Alexa Fluor azide 594 (Invitrogen C10639), according to the manufacturer’s instructions. Samples were then extensively washed with PBS and mounted on glass slides with ProLong Diamond Antifade Mountant (Invitrogen P36961).

### Image acquisition

Images of Figures 2, 3 and 4B were acquired on a Leica TCS SP5 gated-STED microscope, using an HCX PL APO 100X 100/1.40/0.70 oil immersion objective lens (Leica Microsystems, Mannheim, Germany). Emission depletion was accomplished with a 592 nm STED laser. Excitation was provided by a white laser at the desired wavelength for each sample. Alexa488 was excited at 488 nm and its fluorescence emission detected at 500-560 nm, with 1.5-5 ns time gating using a hybrid detector (Leica Microsystems). 512 × 512 pixel images were acquired with a pixel size of 20 nm.

For the experiment reported in Figure 3, the first four STED images were acquired with 16 averages per pixel line, while the fifth image was acquired with 64 averages. We designed this experiment intending to mimic an acquisition with 128 line averages and to be able to monitor the trend of the resolution, noise, and brightness after each 16-averages step. Besides, before and after each STED image, we also acquire a confocal image, in order to monitor the trend of the photobleaching after each STED acquisition. To do so we carefully choose the confocal acquisition parameters in order to induce a confocal-related negligible photobleaching. The final images were then obtained by combining and averaging STED images after each acquisition step so that the resulting image had 16+16 averages, 16+16+16 averages and so on (Figure 3A).

Images of Figure 4G were acquired on a Leica Stellaris 8 Tau-STED microscope, using an HC PL APO CS2 100x/1.40 oil immersion objective lens (Leica Microsystems, Mannheim, Germany). Emission depletion was accomplished with a 775 nm STED laser. Excitation was provided by a white light laser at the desired wavelength for each sample. Alexa594 was excited at 561 nm and its fluorescence emission detected at 570-620 nm using a hybrid detector (Leica Microsystems). 1024 × 1024 pixel images were acquired with a pixel size of 14 nm.

### Generation of SPLIT images

Separation of photon by lifetime tuning (SPLIT) images in Fig.4B were generated using the method of tunable depletion power ^25, 26^. A simplified version of the algorithm described in ^24^ was implemented in Matlab and applied to two-frame stacks consisting of a confocal and a STED image.

### QuICS algorithm

The QuICS analysis was performed in MATLAB (The MathWorks) using a custom code. Given an image I(x,y), a two-dimensional (2D) image correlation function was calculated as:

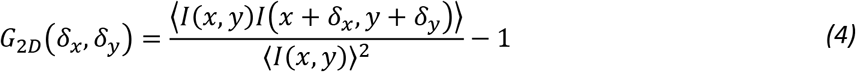

Where the angle brackets indicate averaging over all the selected pixels of the image. The numerator in Eq. (4) was calculated by a 2D fast Fourier transform algorithm. Before calculation, a region of interest (ROI) corresponding to the nucleus was defined using the counterstain signal and the corresponding mask has been applied to the image as described previously ^18, 38^. This step is useful to minimize the effects of nuclear borders on the correlation functions. The 2D correlation function was then converted into one-dimensional radial correlation function, G(ρ), by performing an angular mean ^17^.

To estimate the noise-free correlation function from a single image, we performed a Gaussian fit of the radial correlation function G(ρ) by skipping the first points:

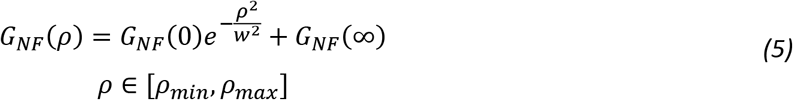

Where the width parameter w corresponds to the 1/e^2^ of a Gaussian function and it is related to the Full Width Half Maximum (FWHM) by the relationship w=FWHM/(2ln2)^1/2^; G_NF_(0) represents the amplitude; G_NF_(∞) represents an offset value. The fitting range was determined as follows. The values ρ_min_ and ρ_max_ were set, by visual inspection of the data, in such a way to exclude the first points and fit a single Gaussian component (Supplementary Figure 1).

As an alternative approach, we generated two independent images I’(x,y) and I’’(x,y) by downsampling the image I(x,y) to half the size. The image I’(x,y) was obtained by averaging the intensity of pixel (i,j) with that of pixel (i+1, j+1), with i+j even. The image I’’(x,y) was obtained by averaging the intensity of pixel (i,j) with that of pixel (i+1, j+1), with i+j odd. The images I’(x,y) and I’’(x,y) were then resampled back to the original size. The 2D cross-correlation function was calculated as:

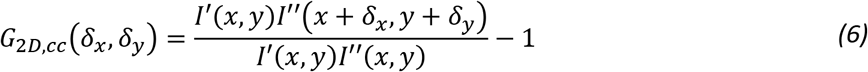

The 2D cross-correlation function was then converted into a one-dimensional radial cross-correlation function, G_cc_(ρ) and fitted with Eq. (5) by setting ρ_min_=0. The two approaches yielded similar results in our data (Supplementary Figure 1).

A user-friendly version of the Matlab code is available at https://github.com/llanzano/QuICS.

## ACKNOWLEDGEMENTS

This work was supported by Associazione Italiana per la Ricerca sul Cancro (AIRC) through MFAG (My First AIRC Grant) Grant ID 21931 and from University of Catania under the program Programma Ricerca di Ateneo UNICT 2020-2022-linea 2. The authors gratefully acknowledge the Bio-Nanotech Research and Innovation Tower (BRIT; PON project financed by the Italian Ministry for Education, University and Research MIUR).

## AUTHOR CONTRIBUTIONS

E.C., A.D. and L.L. designed the study, conceived the experiments and wrote the manuscript. E.C., I.C. and M.D. prepared samples. E.C. and P.B. collected data. L.L. wrote software. E.C., G.I.D, M.F., G.V., P.G.P., A.D. and L.L. analysed data and discussed results. All authors critically reviewed the manuscript.

## ADDITIONAL INFORMATION

The authors declare no competing interests.

---

**Supplementary Figure 1.**
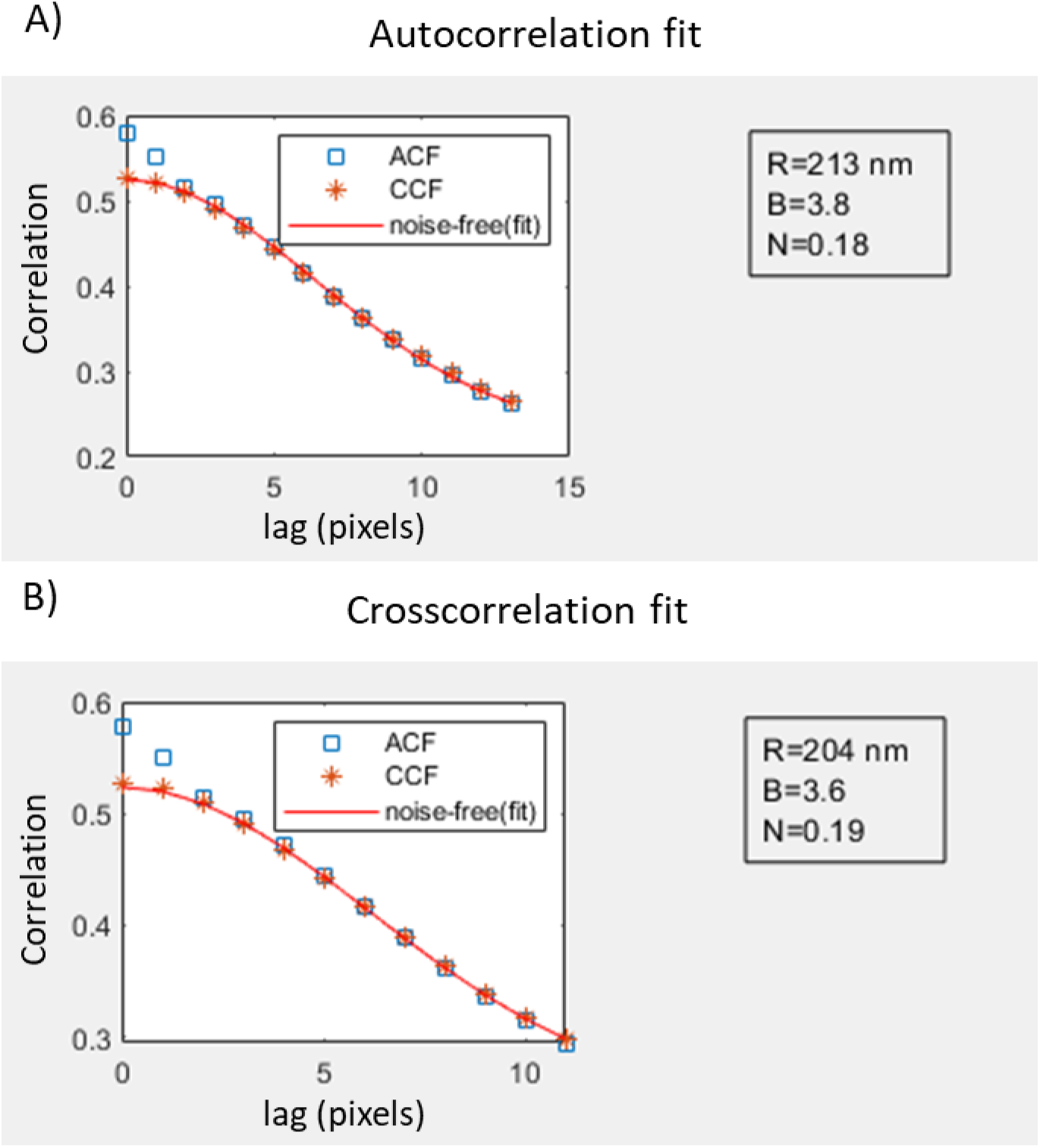
Noise-free correlation function extraction. Shown is an example of data obtained with the QuICS algorithm from the same image. Panel **A)** shows a fit of the autocorrelation function, excluding the first three points, and the relative extracted parameters: R, B, N. Panel **B)** shows the fit of the crosscorrelation function between two statistically independent images obtained through chessboard downsampling, and the relative extracted parameters.

